# The *Rhodobacter sphaeroides* methionine sulfoxide reductase MsrP can reduce *R*- and *S*-diastereomers of methionine sulfoxide from a broad-spectrum of protein substrates

**DOI:** 10.1101/243626

**Authors:** Lionel Tarrago, Sandrine Grosse, Marina I. Siponen, David Lemaire, Béatrice Alonso, Guylaine Miotello, Jean Armengaud, Pascal Arnoux, David Pignol, Monique Sabaty

## Abstract

Methionine (Met) is prone to oxidation and can be converted to Met sulfoxide (MetO), which exists as R- and S-diastereomers. MetO can be reduced back to Met by the ubiquitous methionine sulfoxide reductase (Msr) enzymes. Canonical MsrA and MsrB were shown as absolutely stereospecific for the reduction of S- and R-diastereomer, respectively. Recently, the molybdenum-containing protein MsrP, conserved in all gram-negative bacteria, was shown to be able to reduce MetO of periplasmic proteins without apparent stereospecificity in *Escherichia coli.* Here, we describe the substrate specificity of the *Rhodobacter sphaeroides* MsrP. Proteomics analysis coupled to enzymology approaches indicate that it reduces a broad spectrum of periplasmic oxidized proteins. Moreover, using model proteins, we demonstrated that RsMsrP preferentially reduces unfolded oxidized proteins and we confirmed that this enzyme, like its *E. coli* homolog, can reduce both *R-* and *S*-diastereomers of MetO with similar efficiency.

## Introduction

Aerobic life exposes organisms to reactive oxygen species (ROS) derived from molecular oxygen, such as hydrogen peroxide (H_2_O_2_) or singlet oxygen (^1^O_2_). Bioenergetics chains are important sources of intracellular ROS, with H_2_O_2_ principally produced during respiration (Messner and Imlay, 1999) and ^1^O_2_ arising from photosynthesis (Glaeser et al., 2011). These oxidative molecules act as signaling messengers playing major roles in numerous physiological and pathological states in most organisms and their production and elimination are tightly regulated (Ezraty et al., 2017). However, numerous stresses can affect ROS homeostasis and increase their intracellular concentration up to excessive values leading to uncontrolled reactions with sensitive macromolecules (Imlay, 2013). For instance, photosynthetic organisms, such as plants or the purple bacteria *Rhodobacter sphaeroides* can experience photo-oxidative stress in which unbalance between incident photons and electron transfer in photosynthesis generates detrimental accumulation of ^1^O_2_ (Ziegelhoffer and Donohue, 2009). Moreover, production of ROS could be used advantageously in a defensive strategy against potential pathogenic invaders. For instance, neutrophils produce the strong oxidant hypochlorite (ClO^−^) from H_2_O_2_ and chlorine ions to eliminate bacteria and fungi (Ezraty et al., 2017). Because of their abundance in cells, proteins are the main targets of oxidation (Davies, 2005). Methionine (Met) is particularly prone to oxidation and the reaction of Met with oxidant leads to the formation of Met sulfoxide (MetO), which exists as two diastereomers *R* (Met-*R*-O) and *S* (Met-*S*-O), and further oxidation can form Met sulfone (MetO_2_) (Sharov et al., 1999; Vogt, 1995). Contrary to most oxidative modifications on amino acids, the formation of MetO is reversible, and oxidized proteins can be repaired thanks to methionine sulfoxide reductases (Msr) enzymes that exist principally in two types, MsrA and MsrB. These enzymes, present in almost all organisms, did not evolve from a common ancestor gene and possess an absolute stereospecificity toward their substrates. Indeed, MsrA can reduce only Met-*S*-O (Ejiri et al., 1979; Lowther et al., 2002; Moskovitz et al., 2002; Sharov et al., 1999; Vieira Dos Santos et al., 2005) whereas MsrB acts only on Met-*R*-O (Grimaud et al., 2001; Kumar et al., 2002; Lowther et al., 2002; Moskovitz et al., 2002; Vieira Dos Santos et al., 2005). This strict stereospecificity was enzymatically demonstrated using Met-*R*-O and Met-*S*-O chemically prepared from racemic mixtures of free MetO or using HPLC methods allowing discrimination of both diastereomers, and was structurally explained by deciphering the mirrored pictures of their active site, in which only one MetO diastereomer can be accommodated (Lowther et al., 2002). While MsrA can reduce Met-*S*-O, whether as a free amino acid or included in proteins, MsrB is specialized in the reduction of protein-bound Met-*R*-O, and both are more efficient on unfolded oxidized proteins (Tarrago et al., 2012; Tarrago and Gladyshev, 2012). Eukaryotic Msrs are important actors in oxidative stress protection, aging and neurodegenerative diseases in animals (Kim, 2013), and during environmental stresses or seed longevity in plants (Châtelain et al., 2013; Laugier et al., 2010). In bacteria, MsrA and MsrB are generally located in the cytoplasm (Ezraty et al., 2017), except for *Neisseria* or *Streptococcus* species, for which MsrA and MsrB enzymes can be addressed to the envelope (Saleh et al., 2013; Skaar et al., 2002). They play a role in the protection against oxidative stress and as virulence factors (Ezraty et al., 2017).

Beside these stereotypical MSRs found in all kind of organisms, several bacterial molybdenum cofactor-containing enzymes can reduce MetO. Particularly, the biotin sulfoxide reductase BisC, or its homolog TorZ/BisZ, specifically reduce the free form of Met-*S*-O, in *Escherichia coli* cytoplasm (Ezraty et al., 2005) and *Haemophilus influenza* periplasm, respectively (Dhouib et al., 2016). Moreover, *E. coli* DMSO reductase reduce a broad spectrum of substrates, among which MetO (Weiner et al., 1988), and the *R. sphaeroides* homolog was shown as absolutely stereospecific towards *S*-enantiomer of several alkyl aryl sulfoxides (Abo et al., 1995). Finally, another molybdoenzyme, MsrP (formerly known as YedY), was recently identified as a key player of MetO reduction in the periplasm (Gennaris et al., 2015; Melnyk et al., 2015). MsrP was shown to be induced by exposure to the strong oxidant hypochlorite (ClO^−^) and to reduce MetO on several abundant periplasmic proteins in *E. coli* (Gennaris et al., 2015) or on a Met-rich protein in *Azospira suillum* (Melnyk et al., 2015). A most striking feature of the *E. coli* MsrP (EcMsrP) is that, contrary to all known methionine sulfoxide reductases, it seems capable of reducing both Met-*R*-O and Met-*S*-O (Gennaris et al., 2015). The cistron, *msrP,* belongs to an operon together with the cistron encoding the transmembrane protein MsrQ, which is responsible for the electron transfer to MsrP from the respiratory chain. The operon is conserved in the genome of most gram-negative bacteria suggesting that the MsrP/Q system is very likely a key player for the general protection of the bacterial envelop against deleterious protein oxidation (Gennaris et al., 2015; Melnyk et al., 2015). *R. sphaeroides* MsrP (RsMsrP) shares 50% identical amino acids residues with EcMsrP and transcriptomic analyses evidenced that *RsmsrP* is strongly induced under high-light conditions, suggesting a putative role in protecting the periplasm against ^1^O_2_ (Glaeser et al., 2007).

In this paper, we describe the biochemical characterization of RsMsrP regarding its substrate specificity. Using kinetics activity experiments and mass spectrometry analysis, we show that RsMsrP is a very efficient protein-bound MetO reductase, which lacks stereospecificity and preferentially acts on unfolded oxidized proteins. Proteomics analysis indicate that it can reduce a broad spectrum of proteins in *R. sphaeroides* periplasm, and that Met sensitive to oxidation and efficiently reduced by RsMsrP are found in clusters and in specific amino acids sequences.

## Results

### The *R. sphaeroides* MsrP is an efficient protein-MetO reductase

The results showing that the EcMsrP is a protein-bound MetO reductase, able to reduce both *R-* and *S*-diastereomer of MetO (Gennaris et al., 2015) prompted us to evaluate whether these properties are conserved for RsMsrP. As the EcMsrP was determined to be 5-fold less efficient to reduce the Met-*S*-O than Met-*R*-O, and knowing that all previously identified MetO reductases were absolutely stereospecific toward one enantiomer, we thought that it cannot be excluded that a protein contamination might explain the apparent ability of the EcMsrP to reduce the Met-*S*-O (Gennaris et al., 2015). Such potential Met-*S*-O reductase contaminant should be able to use benzyl viologen (BV) as electron provider and one good candidate is the periplasmic DMSO reductase (Abo et al., 1995; Weiner et al., 1988). Thus, we prepared the recombinant RsMsrP from a *R. sphaeroides* strain devoid of the *dorA* gene encoding the catalytic subunit of the DMSO reductase (Sabaty et al., 2013). After purification on Ni-affinity column and removal of the polyhistidine tag, the mature enzyme was purified by gel filtration, then by strong anion exchange, yielding a highly pure enzyme (Figure S1).

After optimal pH determination (Figure S2), we determined the kinetics parameters of RsMsrP using BV as electron provider and several model substrates: the free amino acid MetO, a synthetic tripeptide Ser-MetO-Ser and the oxidized bovine β-casein (Table 1). The β-casein contains 6 Met, it is intrinsically disordered, and was shown as efficient substrate for the yeast MsrA and MsrB, after oxidation (Tarrago et al., 2012), see also Figure S3). Commercial β-casein contains a mixture of genetic variants, appearing as multiple peaks on mass spectrometry (MS) spectra (Figure 1A). After oxidation with H_2_O_2_, MS analysis confirmed an increase in mass of 96 Da for each peak, very likely corresponding to the addition of 6 oxygen atoms on the Met residues (Figure 1B). Using the free MetO, we determined a *k_cat_* of ~ 122 s^−1^ and a *K_M_* of ~ 115,000 μM, yielding a catalytic efficiency (*k_cat_/K_M_*) of ~ 1,000 M^−1^.s^−1^ (Table 1). With the Ser-MetO-Ser peptide, the *k_cat_* and the *K_M_* values were ~ 108 s^−1^ and ~ 13,000 μM, and thus the *k_cat_/K_M_* was ~ 8,300 M^−1^.s^−1^. Compared to the free MetO, the ~ 8-fold increase in catalytic efficiency is due to the lower *K_M_*, and thus this indicates that the involvement of the MetO in peptide bonds increases its ability to be reduced by the RsMsrP. With the oxidized β-casein, the *k_cat_* and the *K_M_* were ~ 100 s^−1^ and ~ 90 μM, respectively. The *k_cat_/K_M_* was thus ~ 1,000,000 M^−1^.s^−1^. This value, 4 orders of magnitude higher than the one determined with the free MetO, indicates that the oxidized protein is a far better substrate for the RsMsrP. Moreover, even assuming that all MetO in the oxidized β-casein were equal substrates for the RsMsrP and thus multiplying the *K_M_* by 6, the catalytic efficiency obtained (~ 175,000 M^−1^.s^−1^) remained ~ 175-fold higher for the oxidized protein than for the free amino acid. These results indicated that the RsMsrP acts effectively as a protein-MetO reductase.

**Figure 1.**
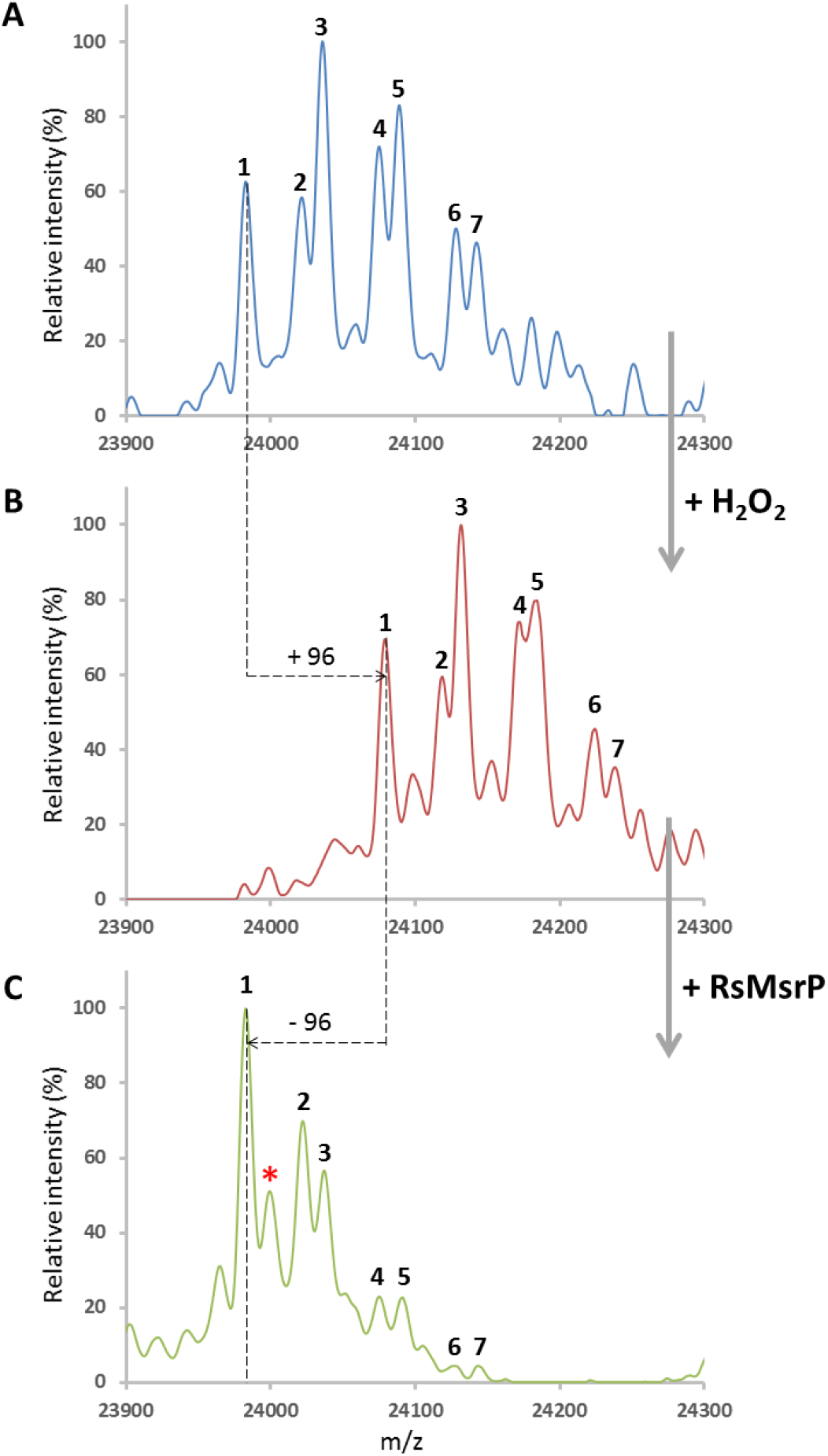
Mass spectrometry spectrum of β-casein non-oxidized (A), oxidized with H_2_O_2_ (B) and repaired by RsMsrP (C). A) Commercial β-casein exists as mixture of genetic variants (7 in our batch). β-casein was analyzed by ESI-MS. Main peaks masses: 1, 23982.7 Da; 2, 24021.6 Da; 3, 24035.9 Da; 4, 24075.0 Da; 5, 24089.1 Da; 6, 24127.5 Da; 7, 24142.5 Da. B) β-casein was oxidized with 50 mM H_2_O_2_ before MS analysis. All major peaks undergone an increase of ~ 96 Da compared to the non-oxidized sample. Main peaks masses: 1, 24079.1 Da; 2, 24118.5 Da; 3, 24131.8 Da; 4, 24172.2 Da; 5, 24184.3 Da; 6, 24224.4 Da; 7, 24238.2 Da. C) Oxidized β-casein was incubated with RsMsrP (25 nM) in presence of BV (0.8 mM) and sodium dithionite (2 mM) as electron donors. All major peaks had masses corresponding of the non-oxidized β-casein, showing the ability to reduce all MetO in this protein. Note the presence of a peak with an increase of 16 Da (*, mass of 23999,4 Da) compared to the main reduced peak, indicating an incomplete reduction of the total protein pool. Main peaks masses: 1, 23983.0 Da; 2, 24022.4 Da; 3, 24037.1 Da; 4, 24075.0 Da; 5, 24091.0 Da; 6, 24128.2 Da; 7, 24143.7 Da.

**Table 1.** Kinetics parameters of RsMsrP reductase activity towards various MetO-containing substrates.

| Substrates | $k_{cat}$ ($s^{-1}$) | $K_M$<br>($\mu M$) | $k_{cat}/K_M$<br>($M^{-1}.s^{-1}$) |
| --- | --- | --- | --- |
| DMSO<sup>a</sup> | $28 \pm 1$ | $61,000 \pm 7,000$ | 465 |
| Free L-Met-*R,S*-O | $122 \pm 20$ | $115,000 \pm 27,000$ | 1,000 |
| Ser-MetO-Ser | $108 \pm 17$ | $13,000 \pm 3,400$ | 8,300 |
| QWGAGM(O)QAEED | $479 \pm 24$ | $4,530 \pm 370$ | 105,700 |
| TTPGYM(O)EEWNK | $> 70$ | $> 5,000$ | N.D. |
| Oxidized β-casein | $100 \pm 5$ | $93 \pm 9$ | 1,075,000 |
| β-casein-*R*-O | $49 \pm 3$ | $51 \pm 6$ | 950,000 |
| β-casein-*S*-O | $8 \pm 1$ | $53 \pm 10$ | 142,000 |
| Oxidized lysozyme | $4 \pm 1$ | $886 \pm 349$ | 4,000 |
| Unfolded oxidized lysozyme | $7 \pm 1$ | $105 \pm 17$ | 70,200 |
| Oxidized GST | $8 \pm 2$ | $643 \pm 194$ | 12,400 |
| Unfolded oxidized GST | $12 \pm 3$ | $99 \pm 33$ | 120,000 |

<sup>a</sup> From Sabaty et al., 2013. *N.D.*, not determined.

### RsMsrP reduces both Met-*R*-O and Met-*S*-O of an oxidized model protein

To determine whether the RsMsrP can reduce both MetO diastereomers, we chose the oxidized bovine β-casein as model substrate because it was efficiently reduced by the yeast MsrA and MsrB indicating the presence of both *R* and *S* diastereomers of MetO (Tarrago et al., 2012). After oxidation with H_2_O_2_, we treated the protein with the MsrA and MsrB, taking advantage of their stereospecificity, to obtain protein samples containing only the Met-*R*-O *(“β-casein-R-O”)* or the Met-*S*-O (“*β-casein-S-O*”), respectively. The absence of one or the other diastereomer of MetO was validated by the absence of remaining Msr activity (Figure S3). These three forms containing two or only one diastereomer of MetO were tested as substrate for RsMsrP (Figure 2). We measured a *k_cat_* of ~ 45 s^−1^ with the oxidized β-casein, which decreased to ~ 30 and to 5 s^−1^ for the β-casein containing the *R* or the *S* sulfoxide, respectively. This result showed that the RsMsrP can reduce both diastereomers of MetO, but appeared 6-fold less efficient to reduce the Met-*S*-O than the Met-*R*-O.

**Figure 2.**
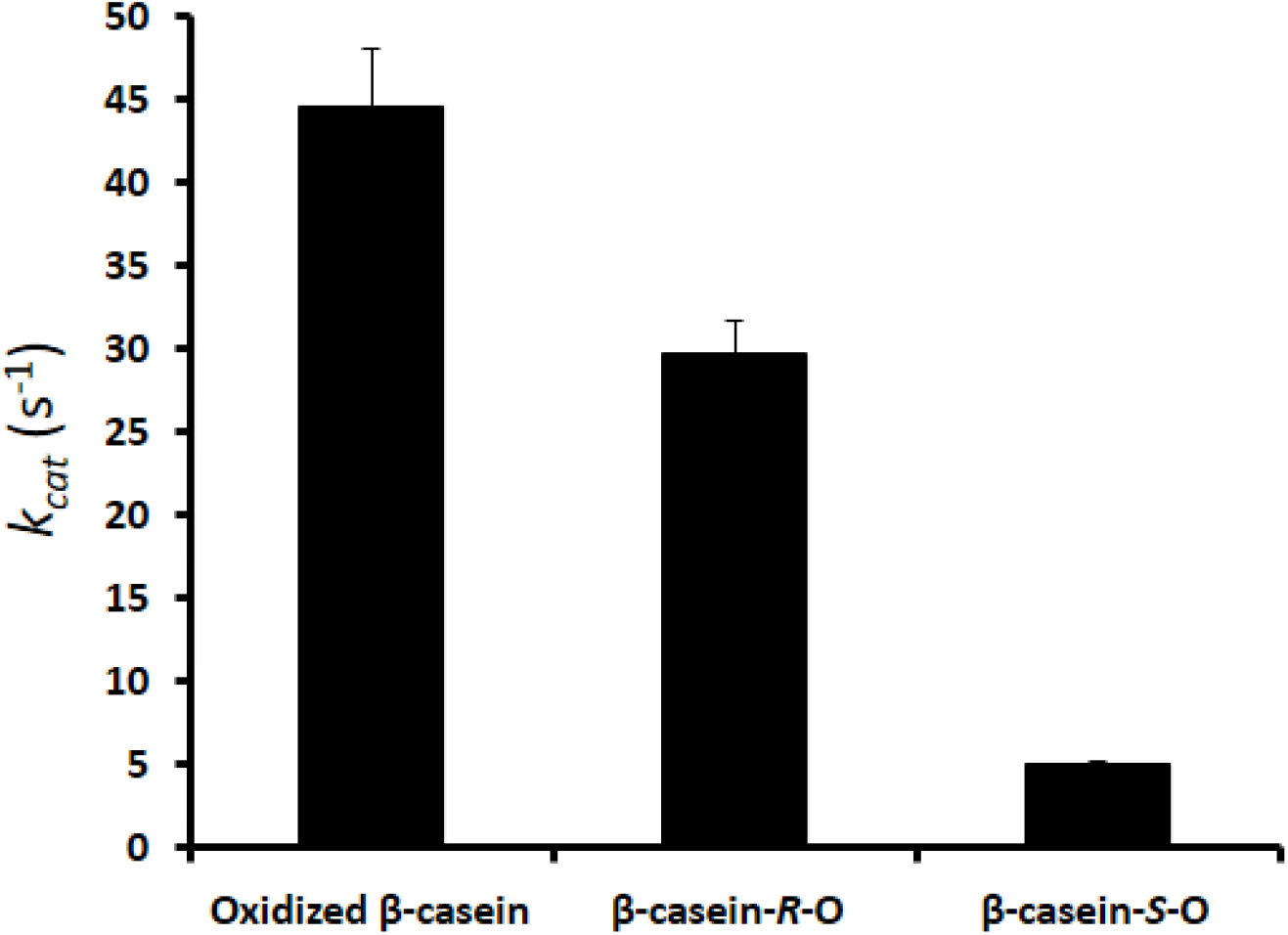
RsMsrP activity using oxidized β-casein, β-casein-*R*-O and β-casein-*S*-O as substrates. The oxidized β-casein (100 μM) containing both diastereomers of MetO, only the *R* one (“*β-casein-R-O*”), or only the *S* one (“*β-casein-S-O*”) were assayed as substrate of RsMsrP. Data presented are average of 3 replicates. ± S.D.

From this result, we postulated that the RsMsrP should be able to reduce all MetO in the oxidized β-casein, as this protein was intrinsically disordered and thus all MetO were very likely accessible. We evaluated this hypothesis by mass spectrometry analysis. When incubated with the RsMsrP, the mass of the oxidized protein decreased of 96 Da, showing that all MetO were reduced (Figure 1C). Altogether, these results clearly showed that the RsMsrP was able to reduce both *R-* and S-diastereomers of MetO contained in the oxidized β-casein, and thus lacked stereospecificity.

### The RsMsrP preferentially reduces Met-*R*-O but acts effectively on Met-*S*-O too

To gain insight into the substrate preference of RsMsrP toward one of the diastereomers of MetO, we performed kinetics analysis using the oxidized β-casein containing the *R* or *S* diastereomers of MetO (Table 1; Figure S4). With the protein containing only the R-diastereomer of MetO (“*β-casein-R-O*”), we determined a *k_cat_* of ~ 50 s^−1^, a *K_M_* of ~ 50 μM and thus a catalytic efficiency of ~ 950,000 M^−1^.s^−1^. In the case of the protein containing only the Met-*S*-O (“*β-casein-S-O*”), the *k_cat_* and *K_M_* were ~ 8 s^−1^ and of ~ 50 μM, respectively. This yielded a catalytic efficiency of 142,000 M^−1^.s^−1^. This value, ~ 7 fold lower than the one obtained with the β-casein-R-O, was due to the lower *k_cat_* as the *K_M_* was not changed. These values seem to indicate that the RsMsrP preferentially reduced the *R* than the *S* diastereomer of MetO in the oxidized β-casein. However, as we could not exclude that the proportion of Met-*R*-O was higher than the proportion of Met-*S*-O in the protein, we developed an assay to estimate the number of MetO reduced by the RsMsrP in the three forms of oxidized β-casein. We measured the total moles of BV consumed for the reduction of all MetO using subsaturating concentrations of the oxidized protein. Practically, the absorbance at 600 nm was measured before and 90 min after the addition of the substrate. As two moles of BV are consumed per mole of MetO reduced, we obtained the apparent stoichiometry of RsMsrP toward the oxidized protein by determining the slope of the linear regression of the straight defined by the amount of MetO reduced as a function of substrate concentration (Figure S5). The values determined were ~ 4.6, ~ 3.2 and ~ 1.8 for the oxidized β-casein, the β-casein-R-O and the β-casein-S-O, respectively. In the case of the oxidized β-casein, we expected a value of 6 based on the data obtained by mass spectrometry (Figure 1). This may have been due to the heterogeneity of the oxidized β-casein (all Met were not fully oxidized initially) and/or to a too short time of incubation (all MetO were not fully reduced, as indicated by the presence of a peak corresponding to a portion of β-casein not fully reduced in Figure 1C). To compare the catalytic parameters, the data were normalized by multiplying the *K_M_* by these apparent stoichiometries, yielding values per MetO reduced and thus allowing the removal of variation due to the different numbers of Met-*R*-O or Met-*S*-O reduced. The catalytic efficiencies were thus 230,000, 300,000 and 80,000 M^−1^.s^−1^ for the oxidized β-casein, the β-casein-*R*-O and the β-casein-*S*-O, respectively (Table 1). The highest value was thus those obtained for the β-casein containing only the *R* form of MetO, indicating that this diastereomer was the preferred substrate for the RsMsrP. However, the value obtained with the β-casein-S-O was only less than 4-fold lower, showing that the RsMsrP can also act effectively on the Met-*S*-O.

### RsMsrP can reduce a broad spectrum of periplasmic proteins

To identify potential periplasmic substrates of RsMsrP and gain insight into its substrate specificity, we applied a high-throughput shotgun proteomic strategy. Periplasmic proteins from *msrP^−^ R. sphaeroides* mutant were extracted, oxidized with NaOCl and then reduced *in vitro* with the recombinant RsMsrP. Untreated periplasmic proteins, oxidized periplasmic proteins and RsMsrP-treated oxidized periplasmic proteins were analyzed by semi-quantitative nanoLC-MS/MS. All experiments were done systematically for 3 biological replicates and resulted in the identification of 362,700 peptide-to-spectrum matches. From all the 11,320 individual peptide sequences, we identified 2,553 unique Met belonging to 720 proteins. The overall percentage of Met oxidation were ~ 35%, ~ 71% and ~ 40% for proteins from the periplasm extract, the oxidized periplasm extract and the RsMsrP-repaired proteins, respectively (Table S1). This first result indicates that the RsMsrP is very likely able to reduce MetO from numerous proteins and to restore an oxidation rate similar to that of the periplasmic extract that has not undergone any oxidation.

The identification of preferential RsMsrP substrates requires the precise comparison of the oxidation state of Met residues from periplasmic proteins before and after the action of the enzyme. After tryptic digestion, since most of the Met/MetO-containing peptides were found in low abundance (i.e. with very low spectral counts), we focused on the proteins robustly detected in all samples. We selected the Met-containing peptides for which at least 10 spectral counts were detected in two replicates for each condition (*i.e.* untreated periplasm, oxidized periplasm and repaired oxidized periplasm) and at least 7 spectral counts were found in the third replicate. This restricted the dataset to 202 unique Met belonging to 70 proteins (Table S2). Overall percentage of Met oxidation (calculated as the number of spectral counts for a MetO-containing peptide vs. the total number of spectral count for this peptide) varied from 2% to 87%, from 9% to 100% and from 4% to 91% in periplasm, oxidized periplasm and repaired oxidized periplasm, respectively. Comparison of Met-O containing peptides between oxidized and RsMsrP treated samples indicates that the percentage of reduction varied from 100 *%* to no reduction at all. 11 MetO were not reduced and 22 were reduced at more than 75 % (only 2 at 100 %). The percentage of reduction for the remaining majority of MetO was almost distributed uniformly between inefficient (less than 25 %) to efficient (75% or more) reduction (Figure 3A).

**Figure 3.**
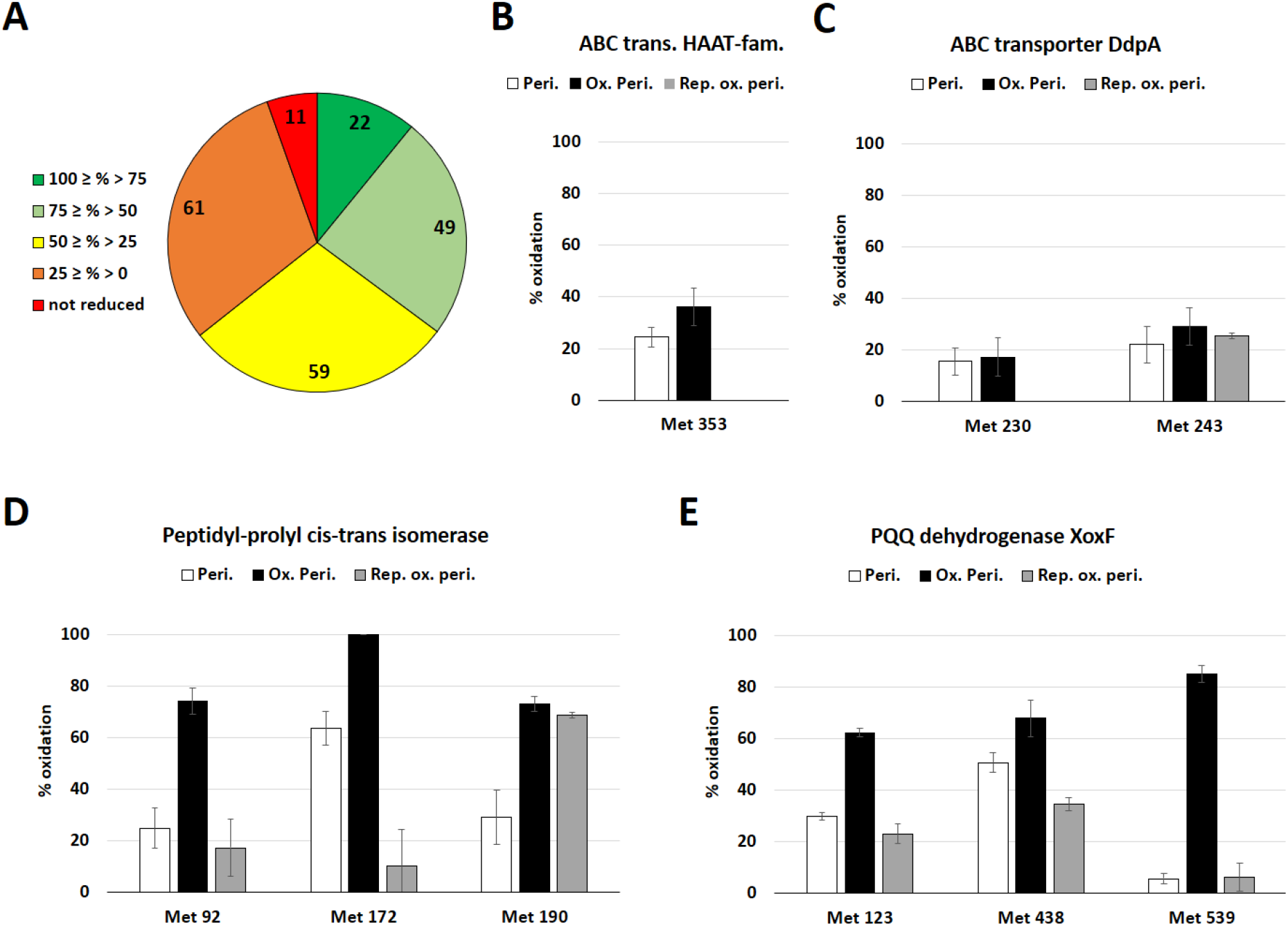
Characteristics of MetO reduction sites and oxidation state of Met in representative proteins. A) Repartition of the number of MetO per percentage of reduction by RsMsrP. B) Percentage of oxidation of Met 353 of putative ABC transporter from HAAT family in the 3 analyzed samples. C) Percentage of oxidation of Met 230 and 243 of ABC transporter DdpA. D) Percentage of oxidation of Met 92, 172 and 190 in the peptidyl-prolyl cis-trans isomerase. E) Percentage of oxidation of Met 123, 438 and 539 of the pyrroloquinoline quinone (PQQ) dehydrogenase XoxF.

No clear evidence of sequence or structure characteristic arose from these 70 identified proteins, neither in term of size or in Met content (Table S2). The periplasmic chaperone SurA, the peptidyl-prolyl cis-trans isomerase PpiA, the thiol-disulfide interchange protein DsbA, the spermidine/putrescine-binding periplasmic protein PotD and the ProX protein were previously proposed as potential substrates of the EcMsrP (Gennaris et al., 2015). All these proteins contain at least one MetO among the most efficiently reduced by the RsMsrP (Table S2), indicating that they are potential conserved substrates of MsrP enzymes in *E. coli* and *R. sphaeroides*, and very likely in numerous gram-negative bacteria.

The sensibility to oxidation of the Met belonging to these 70 proteins, and their efficiency of reduction by the RsMsrP show a wide range of variation, from Met highly sensitive to oxidation and efficiently reduced to Met barely sensitive to NaOCl treatment and not reduced by RsMsrP (Table S2). Moreover, this diversity could be visible within a single protein, in which all Met may not be oxidized and reduced uniformly. For instance, the ABC transporter DdpA, along with another putative ABC transporter (Figure 3B and C), contained one of the two only MetO found as fully reduced in the dataset (Met 230 and Met353, respectively), although DdpA also contained the Met 243 that was neither efficiently oxidized or reduced. This is also illustrated by the case of the peptidyl-prolyl cis-trans isomerase, which possessed the Met found to have the higher decrease in oxidation in all the dataset (Met 172) but also a Met almost not reduced by the RsMsrP (Met 190) (Figure 3D). The Met 539 of the PQQ dehydrogenase XoxF illustrates the case in which a Met was highly sensitive to NaOCl-oxidation and very efficiently reduced (Figure 3E). Twenty-one Met were oxidized at 50 *%* or more and reduced by 50 % or more by RsMsrP (Table S2). Altogether, these results show that RsMsrP can reduce a broad spectrum of apparently unrelated proteins (only 11 Met among 202 were not reduced). However, since all MetO were not reduced with similar efficiency, some structural or sequence determinants could drive the ability of MetO to be reduced by the RsMsrP.

### The nature of the amino acids surrounding a MetO influences the RsMsrP efficiency

Having in hands a relatively large dataset of oxidized and reduced Met prompted us to search for consensus sequences that could favor or impair the oxidation of a Met or the reduction of a MetO by the RsMsrP. For all identified Met, we extracted, the surrounding 5 amino acids on the N- and C-terminal sides to obtain an 11-amino acid sequence with the considered Met centered at the 6^th^ position. We then performed an IceLogo analysis aiming to identify whether some residues were enriched or depleted around the target Met. The principle is to compare a ‘positive’ dataset of peptides, to a ‘negative’ one (Colaert et al., 2009). To find potential consensus sequence of oxidation, we first compared all unique MetO-containing peptides from both the untreated and the NaOCl-oxidized periplasmic extracts, our positive dataset, to the theoretical *R. sphaeroides* proteome used as negative dataset. The IceLogo presented in Figure 4A shows that MetO-containing sequences were mainly depleted of His and aromatic or hydrophobic residues (Trp, Phe, Tyr, Leu, Ile) and were mainly enriched of polar or charged amino acids (Asn, Gln, Asp, Glu and Lys). This suggests that Met in a polar environment, as commonly found at the surface of proteins, are very likely more susceptible to oxidation than those located in hydrophobic environments as in the protein core. We then compared all these unique MetO-containing peptides to all the Met-containing peptides from the same samples (Figure 4B), and we observed that principally Trp, along with His, Tyr and Cys, were depleted around the potentially oxidized Met. Strikingly, the only amino acid significantly more abundant around an oxidized Met was another Met in position −2 and +2. These results indicate that oxidation sensitive Met might be found as clusters.

**Figure 4.**
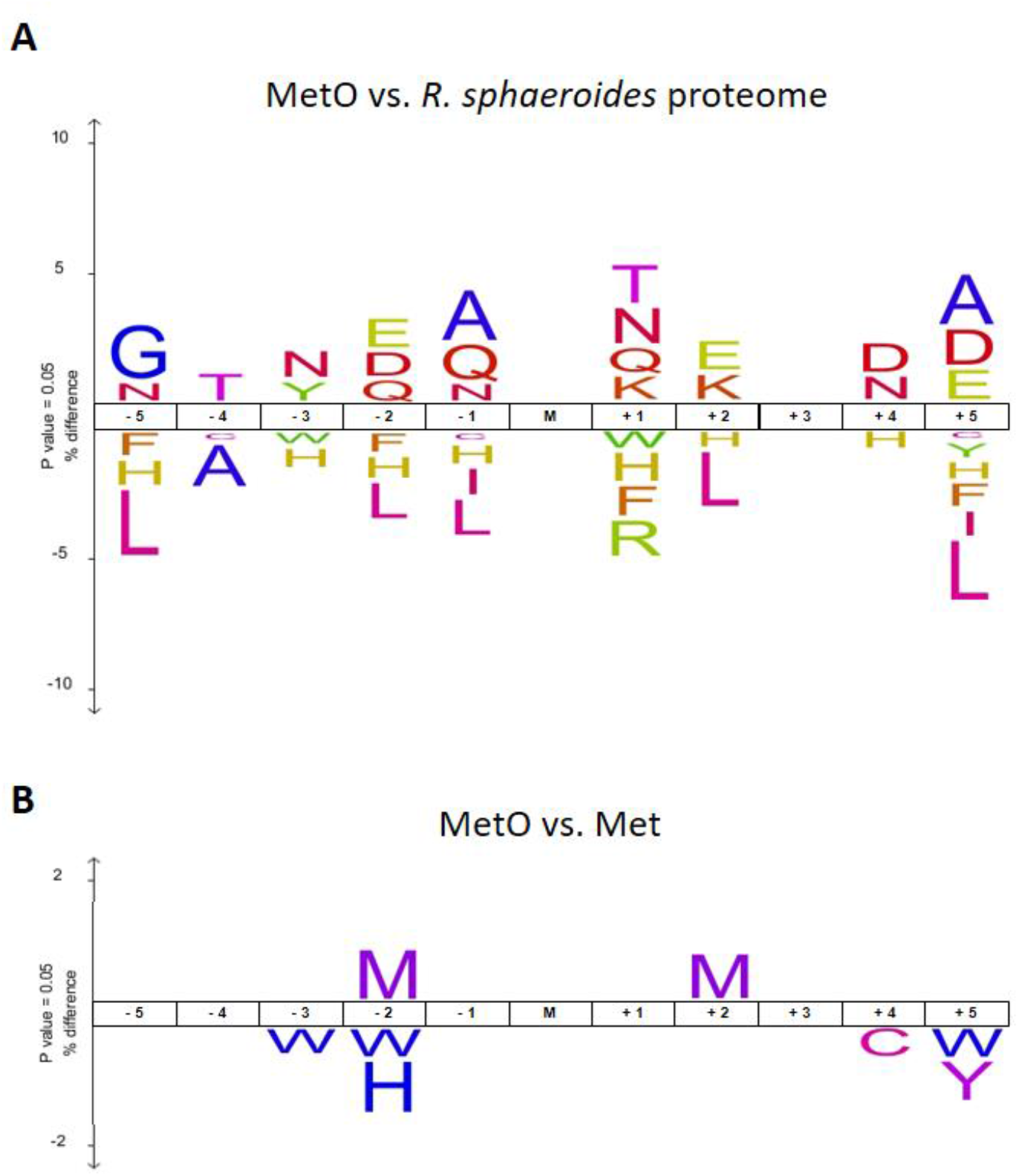
IceLogo representation of enriched and depleted amino acids around site of Met oxidation. A) Enrichment and depletion of amino acids around the oxidized Met (M) found in periplasmic extracts and oxidized periplasmic extracts by comparison with the theoretical proteome of *R. sphaeroides*. B) The same oxidized peptides were analyzed using the peptides containing a non-oxidized Met from the same samples (periplasm and oxidized periplasm extracts).

To identify potential consensus sequence favorable to MetO reduction by the RsMsrP, we performed a precise comparison of the percentage of oxidation before and after the action of the enzyme. We thus defined two criteria to characterize the reduction state of each Met: i) the percentage of reduction calculated using the formula described in Table S2 and based on the comparison of the oxidation percentages in oxidized and repaired oxidized periplasm. For instance, a Met found oxidized at 25 % in the oxidized periplasm and at 5 % in the repaired oxidized periplasm was considered enzymatically reduced at 80 %. ii) the decrease in percentage of oxidation by comparison of the 2 samples. For instance, the same Met found oxidized at 25 % in the oxidized periplasm and 5 % in the repaired extract had a decrease in the percentage of oxidation of 20 %. This second criterion was used to avoid bias in which very little oxidized Met were considered as efficient substrate *(i.e.* a Met oxidized at 5 % in the oxidized periplasm extract and at 1 % in the repaired oxidized periplasm was reduced at 80 %, similarly to one passing from 100 % to 20 %, which intuitively appears as a better substrate than the previous one). We selected as efficiently and inefficiently reduced MetO those for which both criterions were higher than 50 % and lower than 10 %, respectively. The comparison of the sequences surrounding the efficiently reduced MetO to the theoretical proteome of *R. sphaeroides* showed no depletion of amino acid, but mainly enrichment of polar amino acids (Gln, Lys, and Glu) around the oxidized Met (Figure 5A). Similar analysis with the inefficiently reduced MetO indicated the enrichment of Thr and Ser in the far N-terminal positions (−5 and −4) and of a Tyr in position −2 (Figure 5B). The C-terminal positions (+ 1 to +5) were mainly enriched in charged amino acids (Gln, Lys, and Glu), similarly to efficiently reduced MetO. This apparent contradiction may indicate that the amino acids in C-terminal position of the considered MetO did not really influence the efficiency of RsMsrP but were observed simply because of the inherent composition of the overall identified peptides. We then compared the variation of amino acids composition of the MetO-containing peptides between both datasets, using the inefficiently reduced MetO as negative dataset (Figure 5C). The results resembled those obtained by comparison with the entire theoretical proteome of the bacterium, *i.e.* most enriched amino acids were polar (Glu, Gln, Asp and Lys) at most extreme positions (−5, −4 and + 2 to + 5). Of note, the conserved presence of a Gly in position −1, and the presence of several other Met around the central Met. This potential enrichment of Met around an oxidation site is consistent with the result found for the sensibility of oxidation (Figure 4B), and indicates that potential clusters of MetO could be preferred substrates for RsMsrP. We found 16 peptides containing 2 or 3 MetO, reduced at more than 25 % by RsMsrP (Table S2). This was illustrated, for example, by the cell division coordinator CpoB which possesses two close Met residues (66 and 69) highly reduced by the RsMsrP, or by the uncharacterized protein (YP_353998.1) having 4 clusters of MetO reduced by the RsMsrP (Table S2).

From this analysis, the only depleted amino acids appeared to be Thr and Pro in positions −4 and −3 (Figure 4C). To validate these results, we designed two peptides, QWGAGM(O)QAEED and TTPGYM(O)EEWNK, as representative of most efficiently and most inefficiently RsMsrP-reduced peptide-containing MetO, respectively. We used them as substrate to determined kinetics parameters of reduction by RsMsrP (Table 1; Figure S6). The results showed that the peptide QWGAGM(O)QAEED was efficiently reduced, with the highest *k_cat_* value from all the substrates we tested (~ 480 s^−1^) and a *K_M_* of ~ 4,500 μM. This yield a *k_cat_/KM* of ~ 100,000 M^−1^.s^−1^, which is 2 order of magnitude higher than the one determined for the free MetO, and 10-fold lower than for the oxidized β-casein (Table 1). On the contrary, the peptide TTPGYM(O)EEWNK was not efficiently reduced by RsMsrP (Table 1; Figure S6). Indeed, we could not determine the kinetics parameters as activity value curve never reached an inflection point using concentration as high as 5,000 μM. The maximal *k_cat_* value was determined at ~ 70 s^−1^ at 5,000 μM of peptide, which is ~ 3.5-fold less than the one determined with the same concentration of the other peptide (~ 250 s^−1^) (Figure S2). These results are in full agreement with the proteomics analysis and confirm that the nature of the amino acids surrounding a MetO in a peptide or a protein strongly influences its ability to be reduced by RsMsrP.

**Figure 5.**
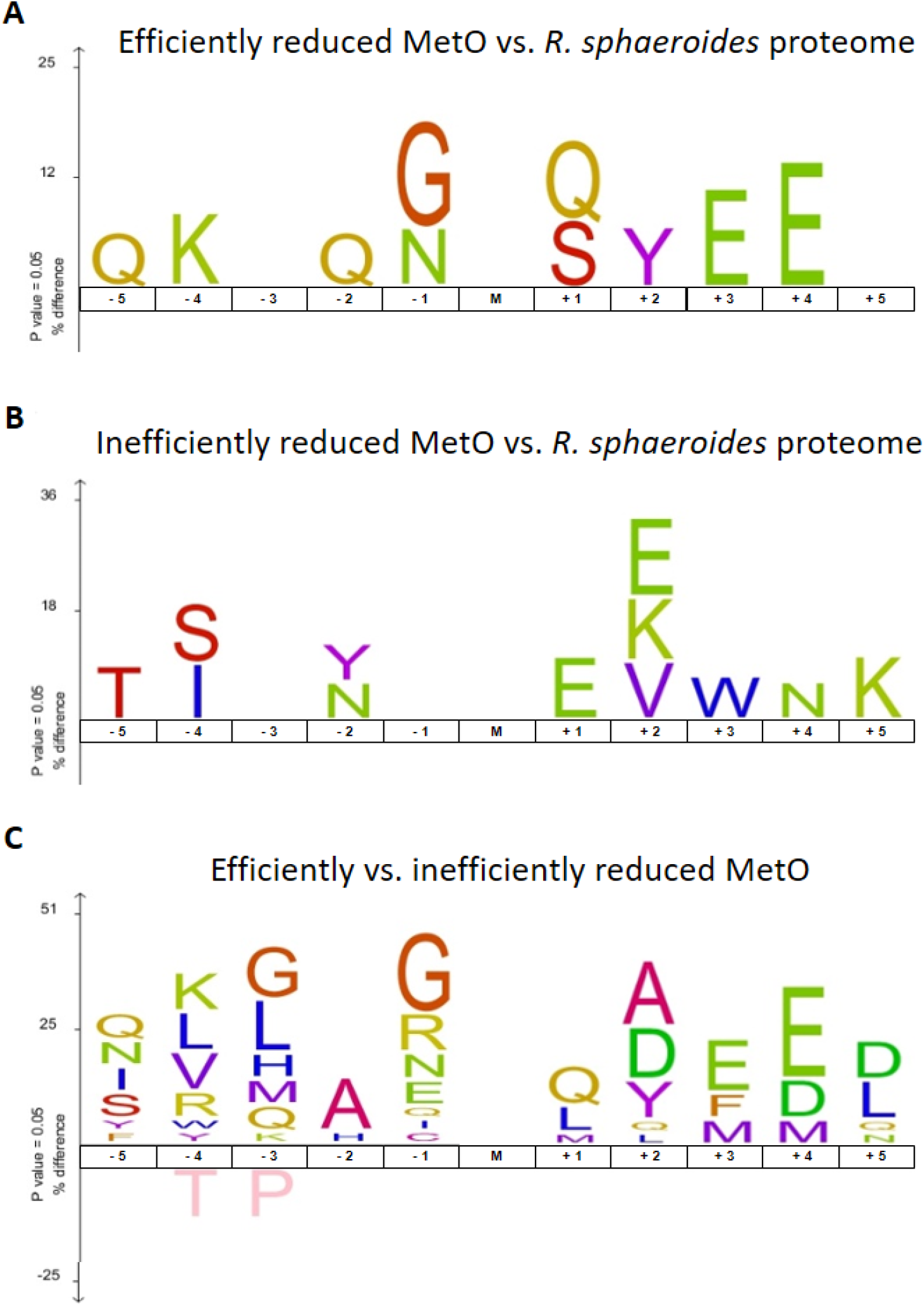
IceLogo representation of enriched and depleted amino acids around site of MetO reduction by RsMsrP. A) Enrichment of amino acids in peptides centered on the MetO for which the percentage of reduction and the decrease in percentage of oxidation were both superior to 50 % by comparison with the theoretical proteome of *R. sphaeroides.* B) Enrichment of amino acids from peptides centered on the MetO for which the percentage of reduction and the decrease in percentage were inferior to 10 % by comparison with the theoretical proteome of *R sphaeroides*. C) Enrichment and depletion of amino acids from efficiently reduced MetO-containing peptides (dataset used in A)) by comparison with inefficiently reduced MetO-containing peptides (dataset used in B)).

### The RsMsrP preferentially reduces unfolded oxidized proteins

To test whether structural determinants affect RsMsrP efficiency of MetO reduction, we compared its activity using oxidized model proteins, either properly folded or unfolded. We started with the chicken lysozyme as it is a very well folded protein highly stabilized with four disulfide bonds (Ray et al., 2001). We oxidized it with H_2_O_2_ and checked its oxidation state by mass spectrometry (Figure S7). Surprisingly, using a protocol similar to the one allowing the complete oxidation of the 6 Met of β-casein, we observed only a weak and incomplete oxidation of the protein. The major peak corresponded to the non-oxidized form and a small fraction had an increase of mass of 16 Da, likely corresponding to the oxidation of one Met. Nevertheless, we prepared from this oxidized sample, an unfolded oxidized lysozyme by reduction with dithiothreitol in 4M urea followed by iodoacetamide alkylation of cysteines, and both samples (oxidized and unfolded oxidized), were used as substrates for RsMsrP (Figure 6). We also used glutathione-*S*-transferase (GST) which possesses 9 Met and is highly structured. After oxidation with H_2_O_2_, GST was incubated with 4 M of the chaotropic agent urea, a concentration sufficient to induce complete unfolding of the protein (Tarrago et al., 2012). For both oxidized proteins, we observed a dramatic increase in activity after unfolding. Indeed, the RsMsrP activity increased 7-fold with the unfolded oxidized lysozyme compared to the folded one, and 6-fold in the case of the unfolded oxidized GST compared to the folded oxidized GST (Figure 6). As the unfolded oxidized protein solutions of lysozyme or GST contained a substantial amount of urea, we made controls in which the urea was added extemporaneously in the cuvette during the measurements, showing that urea did not influence the RsMsrP activity (Figure S8).

**Figure 6.**
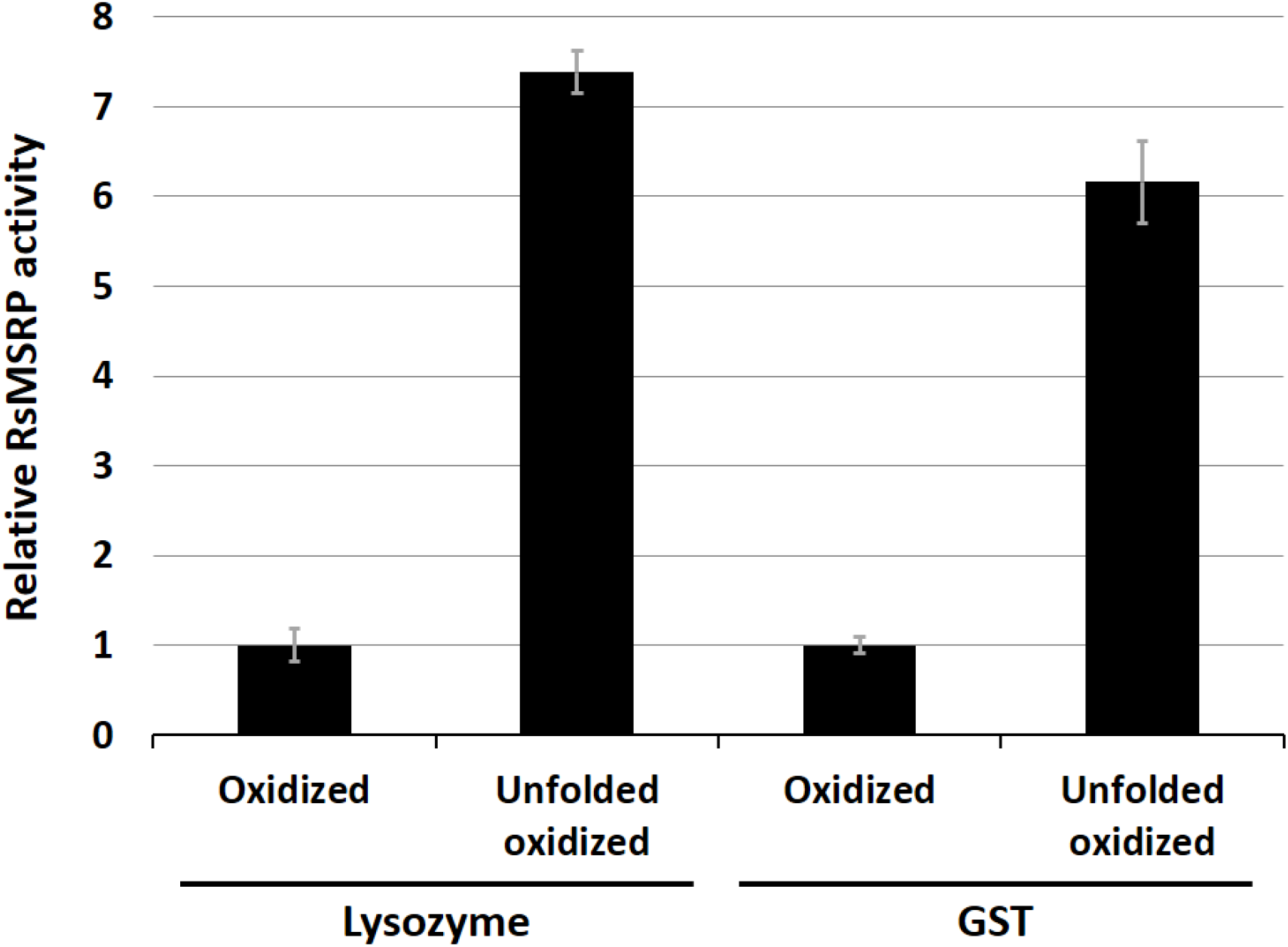
Relative RsMsrP activity using unfolded oxidized proteins. The RsMsrP activity was determined as described in Figure S4. Oxidized and unfolded oxidized lysozyme were incubated at 100 μM in 50 mM MES, pH 6.0. Initial turnover numbers were 0.65 ± 0.12 s^−1^ and 7.38 ± 0.23 s^−1^ with oxidized and unfolded oxidized lysozyme, respectively. Activity with oxidized and unfolded oxidized GST (75 μM) was determined similarly except that reaction buffer was 30 mM Tris-HCl, pH 8.0 because unfolded oxidized GST precipitated in 50 mM MES, pH 6.0. Initial turnover numbers were 0.86 ± 0.08 s^−1^ and 5.31 ± 0.39 s^−1^ with oxidized and unfolded oxidized GST, respectively. Data presented are average of three replicates. ± S.D.

Mass spectrometry analysis showed that the RsMsrP was able to completely reduce the oxidized lysozyme in these conditions (Figure S7), suggesting that observed differences of repair between the folded- and unfolded-oxidized lysozyme were not due to the incapacity of the RsMSRP to reduce some MetO, but were due to kinetics parameters. We thus determined the kinetic of reduction of these proteins by the RsMsrP (Table 1, Figure S8). With the oxidized lysozyme, the *k_cat_* and the *K_M_* were ~ 4 s^−1^ and ~ 900 μM, respectively. Using the unfolded oxidized lysozyme, the *k_cat_* increased to ~ 7 s^−1^ and the *K_M_* decreased to ~ 100 μM. The catalytic efficiency determined with the unfolded oxidized lysozyme was thus ~ 18-fold higher than the one determined using the oxidized lysozyme before unfolding (70,200 vs. 4,000 M^−1^. s^−1^). Similar results were obtained with the GST. Indeed, with the oxidized GST, we recorded *k_cat_* and *K_M_* values of ~ 8 s^−1^ and ~ 640 μM, respectively whereas for the unfolded oxidized GST, the *k_cat_* was slightly higher (~ 12 s^−1^), and the *K_M_* was ~ 6-fold lower (~ 100 μM). The catalytic efficiency was 10-fold higher for the unfolded oxidized GST than for its folded counterpart (Table 1; Figure S8). Altogether, these results showed that the RsMsrP is more efficient in reducing MetO in unfolded than in folded oxidized proteins. Moreover, as evidenced with lysozyme that contained only one MetO in our conditions, the increase in activity using unfolded substrate is not dependent of the number of MetO reduced.

## Discussion

All organisms have to face harmful protein oxidation and almost all possess canonical Msrs that protect proteins by reducing MetO. Bacteria also have molybdoenzymes able to reduce MetO, as a free amino acid for the DMSO reductase (Weiner et al., 1988) or the biotin sulfoxide reductase BisC/Z (Dhouib et al., 2016; Ezraty et al., 2005), but also included in proteins in the case of the MsrP (Gennaris et al., 2015; Melnyk et al., 2015). Genetic studies and the conservation of MsrP in most gram-negative bacteria indicate that it is very likely a key player in the protection of periplasmic proteins against oxidative stress (Gennaris et al., 2015; Melnyk et al., 2015) However, an in-depth characterization of its protein substrate specificity is still lacking. In this work, we chose the MsrP from the photosynthetic purple bacteria *R. sphaeroides* as model enzyme to uncover such specificity. Using purified oxidized proteins and peptides, we showed that RsMsrP is a very efficient protein-containing MetO reductase, with apparent affinities (*K_M_*) for oxidized proteins 10 to 100-fold lower than for the tripeptide Ser-MetO-Ser or the free MetO (Table 1). As reported for canonical MsrA and MsrB (Tarrago et al., 2012), we observed important variations in the *k_cat_* of reduction of different oxidized proteins, arguing for the existence of sequence and structural determinant affecting the enzyme efficiency (Table 1).

To find potential physiological substrates of RsMsrP and uncover their properties, we used a proteomic approach aiming to compare the oxidation state of periplasmic proteins after treatment with the strong oxidant NaOCl, then followed by RsMsrP reduction. We found 202 unique Met, belonging to 70 proteins, for which the sensitivity of oxidation and the ability to serve as RsMsrP substrate varied greatly (Figure 3, Table S2). MetO efficiently reduced by RsMsrP belong to structurally and functionally unrelated proteins, indicating that RsMsrP very likely does not possess specific substrates and acts as a global protector of protein integrity in the periplasm. Interestingly, we observed from our IceLogo analysis that Met sensitive to oxidation are generally presented in a polar amino acid environment and can be found in cluster (Figure 4). These properties might be common to all Met in proteins as similar results were found in human cells (Ghesquière et al., 2011; Hsieh et al., 2017) and plants (Jacques et al., 2015). Moreover, oxidized Met efficiently reduced by the RsMsrP were also found in cluster in polar environment and our analysis shows that the presence of Thr and Pro in N-terminal side of a MetO strongly decrease RsMsrP efficiency (Table 1, Figure 5 and S6). To our knowledge, the presence of a Thr close to a MetO was not previously shown to influence any Msr activity, but the presence of a Pro was shown to decrease or totally inhibit MetO reduction by the human MsrA and MsrB3, depending on its position (Ghesquière et al., 2011).

The presence of oxidation-sensitive Met efficiently reduced by the RsMsrP in clusters on polar parts of proteins should facilitate the oxidation/reduction cycle aiming to scavenge ROS as previously proposed for canonical Msrs (Luo and Levine, 2009). This is also illustrated by the methionine-rich protein MrpX proposed as main substrate of the *A. suillum* MsrP, which is almost only composed of Met, Lys, Glu and Asp (Melnyk et al., 2015). The presence of numerous MetO on a single molecule of protein substrate should increase the RsMsrP efficiency as one molecule of substrate allows several catalytic cycles, potentially without breaking physical contact between the enzyme and its substrate.

Comparison of the RsMsrP activity using folded or unfolded protein substrates (lysozyme and GST) showed that it is far more efficient to reduce unfolded oxidized proteins (Figure 6). Similar results were found for canonical Msrs (Tarrago et al., 2012). In the case of the MsrB it was because more MetO were accessible for reduction whereas for MsrA this increase was independent of the number of MetO reduced. Here, the use of the lysozyme containing only one MetO (Figure S7) undoubtedly showed that the increase in activity is not related to the unmasking of additional MetO upon protein denaturation (Table 1; Figure 6). This could indicate that the RsMsrP has a better access to the MetO in the protein or that the MetO is more easily accommodated in the active site of the enzyme because of increased flexibility. This should provide a physiological advantage to the bacteria during oxidative attacks, which could occur during other stresses such as acid or heat, hence promoting simultaneous oxidation and unfolding of proteins. Particularly, hypochlorous acid, shown to induce *msrP* expression in *E. coli* (Gennaris et al., 2015) and *A. suillum* (Melnyk et al., 2015) has strong oxidative and unfolding effect on target proteins (Winter et al., 2008).

Finally, previous work indicated that the *E. coli* MsrP lacks stereospecificity and can reduce both R- and *S*-diastereomers of MetO chemically isolated from a racemic mixture of free L-Met-R, *S*-O (Gennaris et al., 2015). This discovery is of fundamental importance as it breaks a paradigm in the knowledge about Met oxidation and reduction, and very likely for all enzymology as non-stereospecific enzymes were very rarely described. Indeed, to our knowledge, all previously characterized enzymes able to reduce Met sulfoxide or related substrates were shown as absolutely stereospecific. This was the case for the canonical MsrA and MsrB, which reduce only the *S*-diastereomer and the R-diastereomer, respectively (Ejiri et al., 1979; Grimaud et al., 2001; Kumar et al., 2002; Lowther et al., 2002; Moskovitz et al., 2002; Sharov et al., 1999; Vieira Dos Santos et al., 2005), as well as for the free Met-*R*-O reductase (Le et al., 2009; Lin et al., 2007) and for the molybdoenzymes DMSO reductase (Abo et al., 1995; Weiner et al., 1988) and BisC/Z (Dhouib et al., 2016; Ezraty et al., 2005). To evaluate the potential lack of stereospecificity of the RsMsrP, we chose to use a strategy different than the one used for *E. coli* MsrP (Gennaris et al., 2015) and prepared oxidized β-casein containing only one or the other MetO diastereomer using yeast MsrA and MsrB to eliminate the *S*- and the *R*-diastereomers, respectively. Activity assays and kinetics experiments using a highly purified RsMsrP demonstrated that it can efficiently reduce the β-casein containing only the R- or the *S*-diastereomer (Table 1; Figure 2 and S4). Moreover, this lack of stereospecificity was undoubtedly confirmed by the ability of the RsMsrP to reduce all 6 MetO formed on the oxidized β-casein (Figure 1). These results, consistent with Gennaris and coworkers finding, indicate that this lack of stereospecificity is very likely common to all MsrP homologs. Together with the apparent ability of the enzyme to repair numerous unrelated oxidized proteins, the capacity to reduce both diastereomers of MetO, argues for a role of MsrP in the general protection of envelope integrity in gram negative bacteria. However, it raises questions regarding the structure of its active site as the enzyme should be able to accommodate both diastereomers. From this, we wondered whether the RsMsrP could reduce the Met sulfone, which can be imagined as a form of oxidized Met containing both *R*- and *S*-diastereomers, but we did not detect any activity (Figure S9). Although it could be because of an incompatibility in redox potential, it may indicate that this form of oxidized Met cannot reach the catalytic atom. The three-dimensional structure of the oxidized form of *E. coli* MsrP indicated that the molybdenum atom, which is supposed to be the catalytic center of the enzyme, is buried 16 Å from the surface of the protein (Loschi et al., 2004). The next challenge will be to understand the MsrP reaction mechanism and will require the determination of the enzyme structure in its oxidized and reduced forms bound to its MetO-containing substrates.

## Significance

Protein quality control is a vital cellular process. The cell envelope and the periplasm of gram-negative bacteria are particularly exposed to oxidative molecules damaging proteins. Methionine residues are prone to oxidation, and can be converted to two diastereomers, *R* and *S*, of methionine sulfoxide (MetO). Almost all organisms possess the thiol-oxidoreductases, methionine sulfoxide reductases (Msr) A and B able to reduce the *S*- and R-MetO, respectively, with a strict stereospecificity. Recently, a new enzymatic system, MsrQ/MsrP which is conserved in all gram-negative bacteria, was identified as a key actor in the reduction of oxidized proteins in periplasm. The haem-binding membrane protein MsrQ transmits the reducing power coming from the electron transport chains to the molybdoenzyme MsrP which acts as protein-MetO reductase. MsrQ/MsrP function was genetically well established, but the identity and the biochemical properties of MsrP substrates remain unknown. In this work, using the purified MsrP enzyme from the photosynthetic bacteria *Rhodobacter sphaeroides* as a model, we show that it can reduce a broad spectrum of protein substrates. The most efficiently reduced MetO are found in cluster in amino acids sequence devoid of threonine and proline in C-terminal side. Moreover, *R. sphaeroides* MsrP lacks stereospecificity as it can reduce both *R* and *S* diastereomers of MetO, like its *Escherichia coli* homolog, and preferentially acts on unfolded oxidized proteins. Due to its high conservation of in all gram-negative, understanding how the MsrQ/MsrP system protects periplasmic proteins from oxidation and helps bacteria to cope with harmful environment is of fundamental importance and could provide insight to create molecules to fight pathogenic bacteria.

## Acknowledgments

We are very grateful to Prof. Vadim, N. Gladyshev (Brigham’s and Women Hospital and Harvard Medical School) for the gift of pET28a-MsrA, pET21b-MsrB, pET15b-TR1, pET15b-Trx1 and pGEX4T1 expression vectors. Pascaline Auroy-Tarrago (Laboratoire de Bioénergétique et Biotechnologie des Bactéries et Microalgues, CEA, BIAM) is acknowledged for her help with proteomics analysis. This work was supported by the Commissariat à l’Energie Atomique et aux Energies Alternatives (CEA) and by the project METOXIC (ANR 16-CE11-0012).

## Author contributions

LT, PA, DP and MS designed the study. LT, SG, MIS and MS purified RsMsrP. LT and MS prepared all other proteins. LT, SG, MIS, MS performed biochemical characterization of RsMsrP. LT, MS and DL performed β-casein and lysozyme mass spectrometry analysis and analyzed the data. SG and MS prepared *R. sphaeroides* 2.4.1 *msrP^−^* mutant and periplasmic proteins samples. BA, GM and JA performed proteomics analysis of periplasmic proteins and LT, MS, GM and JA analyzed the data. LT wrote the manuscript with contribution of DL, PA, DP, JA and MS. All authors approved the final manuscript.

## Materials and methods

### Production and purification of recombinant proteins

Recombinant MsrP was produced similarly as described in (Sabaty et al., 2013). Briefly, *R. sphaeroides* f sp. *denitrificans* IL106 *dmsA^−^* strain carrying pSM189 plasmid allowing the production of a periplasmic MsrP with a 6-His N-terminal tag was grown in 6-liter culture under semi-aerobic conditions in Hutner medium until late exponential phase. Periplasmic fraction was extracted and loaded on HisTrap column (GE Healthcare) then MsrP was eluted by an imidazole step gradient. MsrP solution was concentrated using 15-ml Amicon^®^ Ultra concentrators with 10-kDa cutoff (Millipore), desalted with Sephadex G-25 in PD-10 Desalting Columns (GE Healthcare). The protein concentration was adjusted to 1 mg.ml^−1^ in Tris-HCl 30 mM, 500 mM NaCl, pH 7.5, the Tobacco Etch Virus (TEV) protease was added (1:80 TEV:RsMsrP mass ratio) and the solution incubated overnight at room temperature to remove the polyhistidine tag. Untagged RsMsrP was purified on a second HisTrap column, then concentrated and desalted in 50 mM 4-(2-hydroxyethyl)-1-piperazineethanesulfonic acid (HEPES), pH 8.0. Protein solution was then loaded on gel filtration in Superdex™ 200 10/30 column equilibrated with Tris-HCl 30 mM, pH 7.5. Main fractions were pooled and applied to a MonoQ™ 4.6/100 PE (GE Healthcare). RsMsrP was then eluted using a linear NaCl gradient (0 to 500 mM). Fractions were analyzed on SDS-PAGE using NuPAGE™, 10 % Bis-Tris gels with MES-SDS buffer (ThermoFisher). Recombinant MsrA, MsrB, Thioredoxin Reductase (TR) 1, Thioredoxin 1 (Trx1) from *Saccharomyces cerevisiae* and containing a polyhistidine tag, as well as the glutathione-S-transferase (GST) from *Schistosoma japonicum,* were produced and purified as previously described (Tarrago et al., 2012). Protein concentrations were determined spectrophotometrically using specific molar extinction coefficients at 280 nm: 6-His-RsMSRP, 56,380 M^−1^.cm^−1^; untagged RsMsrP, 54,890 M^−1^.cm^−1^; MsrA, 34,630 M^−1^.cm^−1^; MsrB, 24,325 M^−1^.cm^−1^; TR1, 24,410 M^−1^.cm^−1^; Trx1, 9,970 M^−1^.cm^−1^; GST, 42,860 M^−1^.cm^−1^, bovine β-casein (Sigma-Aldrich), 11,460 M^−1^.cm^−1^ and chicken lysozyme (Sigma-Aldrich), 32,300 M^−1^.cm^−1^. Protein solutions were stored at −20°C until further use.

### Peptides

Ser-Met(O)-Ser, QWGAGM(O)QAEED and TTPGYM(O)EEWNK peptides were obtained from GenScript^®^ (Hong-Kong).

### Preparation of oxidized bovine β-casein and its Met-*R*-O and Met-*S*-O containing counterparts

For oxidation, bovine β-casein was incubated in Phosphate Buffer Saline (PBS) at 1 mg.ml^−1^ in the presence of 200 mM H_2_O_2_ and incubated overnight at room temperature. H_2_O_2_ was removed by desalting using PD-10 column and the protein solution was concentrated with 10-kDa cutoff Amicon^®^ Ultra concentrator. Oxidized GST was prepared similarly using 100 mM H_2_O_2_. To prepare Met-*R*-O containing β-casein, a solution of oxidized β-casein was incubated in Tris-HCl 30 mM, pH 8 at a final concentration of 6.5 mg.ml^−1^ (260 μM) in the presence of 25 mM dithiothreitol (DTT) and 10 μM MsrA and incubated overnight at room temperature. The solution was 10-fold diluted in Tris-HCl 30 mM, pH 8 and passed on HisTrap column to remove the his-tagged MsrA. After concentration, the DTT was removed by desalting using PD-10 column. Met-*S*-O containing β-casein was prepared similarly replacing the MsrA by the MsrB (14 μM). The protein solutions were concentrated with 10-kDa cutoff Amicon^^®^^ Ultra concentrator and the final concentration was determined spectrophotometrically. Protein solutions were stored at −20°C until further use.

### Enzymatic activity and apparent stoichiometry measurements

RsMsrP reductase activity was measured as described in(Sabaty et al., 2013) with few modifications. Benzyl viologen was used as electron donor and its consumption was followed at 600 nm using at UVmc1^®^ spectrophotometer (SAFAS Monaco) equipped with optic fibers in a glovebox workstation (MBRAUN Labstar) flushed with nitrogen. We determined the specific molar extinction coefficient of benzyl viologen at 8,700 M^−1^.cm^−1^ in 50 mM 2-(N-morpholino)ethanesulfonic acid (MES), pH 6.0 buffer. Each reaction mixture (1 ml or 0.5 ml) contained 0.2 mM benzyl viologen reduced with sodium dithionite, and variable concentrations of substrates in 50 mM MES, pH 6.0 buffer.

Reactions were started by addition of the RsMsrP enzyme (10 to 46 nM). Reduction of MetO rates were calculated from ΔA_600_ nm slopes respecting a stoichiometry of 2 (2 moles of benzyl viologen are oxidized for 1 mole of MetO reduced).

The apparent stoichiometry was determined similarly, using subsaturating concentrations of substrates: 1–10 μM oxidized β-casein, 1–10 μM Met-*R*-O containing β-casein and 1.5–15 μM Met-*S*-O containing β-casein. The amount of oxidized benzyl viologen was determined 1 hour after the addition of the RsMsrP (46 nM) by subtracting the final A_600_ nm value to the initial one. Controls were made without the RsMsrP enzyme, and without the MetO-containing substrate. Quantities of MetO reduced were plotted as function of substrates quantities and the apparent stoichiometry was obtained from the slope of the linear regression.

MsrA and MsrB activities were measured following the consumption of NADPH spectrophotometrically at 340 nm using the thioredoxin system similarly as previously described (Tarrago et al., 2012). A 500-μl reaction cuvette contained 200 μM NADPH, 2 μM TR1, 25 μM Trx1 and 5 μM MsrA or MsrB and 100 μM oxidized β-casein. Production of Met was calculated respecting a stoichiometry of 1 (1 mole of NADPH is oxidized for 1 mole of Met produced).

Analysis and kinetics parameters determination were made using GraphPad^®^ Prism 4.0 software (La Jolla, CA, USA).

### Electrospray ionization/Mass spectrometry analysis of purified proteins

For oxidation, bovine β-casein (5 mg.ml^−1^) in 50 mM HEPES, pH 7.0, was incubated overnight at room temperature with H_2_O_2_ (50 mM). H_2_O_2_ was removed by desalting using PD-10 column and the protein solution was concentrated with 10-kDa cutoff Amicon^®^ Ultra concentrator. Oxidized β-casein (100 μM) was reduced by addition of 44 nM MsrP in a reaction mixture containing 50 mM HEPES pH 7.0, 0.8 mM benzyl viologen and 0.2 mM sodium dithionite. After two hours reaction in the glove-box, the repaired β-casein was analyzed by mass spectrometry in comparison to non-oxidized and oxidized β-casein.

Mass spectrometry analyses were performed on a MicroTOF-Q Bruker (Wissembourg, France) with an electrospray ionization source. Samples were desalted and concentrated in ammonium acetate buffer (20 mM) (Sigma-Aldrich) prior analyses with Centricon Amicon (Millipore) with a cutoff of 30 kDa. Samples were diluted with CH3CN/H2O (1/1-v/v), 0.2% Formic Acid (Sigma). Samples were continuously infused at a flow rate of 3 μL/min. Mass spectra were recorded in the 50-7000 mass-to-charge (m/z) range. MS experiments were carried out with a capillary voltage set at 4.5 kV and an end plate off set voltage at 500 V. The gas nebulizer (N2) pressure was set at 0.4 bars and the dry gas flow (N2) at 4 L/min at a temperature of 190 °C. Data were acquired in the positive mode and calibration was performed using a calibrating solution of ESI Tune Mix (Agilent) in CH3CN/H2O (95/5-v/v). The system was controlled with the software package MicrOTOF Control 2.2 and data were processed with DataAnalysis 3.4.

### Generation of *R. sphaeroides* 2.4.1 *msrP^−^* mutant

The *msrPQ* operon was amplified from *R. sphaeroides* 2.4.1 genomic DNA with the primers 5’-AGATCGACACGCCATTCACC-3’ and 5’-TCGGTGAGGCGCTATCTAGG-3’. The 2.2 kb PCR product was cloned into pGEMT Easy (Promega). An omega cartridge encoding resistance to streptomycin and spectinomycin (Prentki and Krisch, 1984) was then cloned into the *BamHI* site of *msrP.* The resulting plasmid was digested with *SacI* and the fragment containing the disrupted *msrP* gene was cloned into pJQ200mp18 (Quandt and Hynes, 1993). The obtained plasmid, unable to replicate in *R. sphaeroides*, was transferred from *E. coli* by conjugation. The occurrence of a double-crossing over event was confirmed by PCR and absence of the protein from the SDS-PAGE profile.

### Preparation of periplasmic samples for proteomics analysis

*R. sphaeroides* 2.4.1 *msrp*^−^ mutant was grown under semi-aerobic conditions. Periplasmic extract was prepared as previously described (Sabaty et al., 2010) by cells incubation in 50 mM HEPES pH 8.0, 0.45 M sucrose, 1.3 mM Ethylenediaminetetraacetic acid (EDTA) and 1 mg.ml^−1^ chicken lysozyme. For Met oxidation, the periplasmic extract (0.7 mg.ml^−1^) was incubated with 20 mM *N*-Ethylmaleimide (NEM) and 2 mM NaOCl (Sigma-Aldrich) in 50 mM HEPES pH 8.0, 50 mM NaCl for 10 min at room temperature. NaOCl was removed by desalting using PD-10 column and buffer was changed for 50 mM MES pH 6.0. The protein solution was concentrated with 3-kDa cutoff Amicon^®^ Ultra concentrator. Three reaction mixtures were prepared in the glove box containing 35 μl of periplasmic extract, 1 mM benzyl viologen, 2 mM dithionite in 50 mM MES pH 6.0. The protein concentration in each reaction was 2.5 mg.ml^−1^. The first reaction contained non-oxidized periplasmic extract, the second and third ones contained oxidized periplasmic extract. For the third reaction (repaired periplasm) 10 μM RsMsrP was added. The reactions were incubated for three hours at room temperature.

### Trypsin proteolysis and tandem mass spectrometry

Protein extracts were immediately subjected to denaturing PAGE electrophoresis for 5 min onto a 4–12% gradient 10-well NuPAGE (Invitrogen) gel. The proteins were stained with Coomassie Blue Safe solution (Invitrogen). Polyacrylamide bands corresponding to the whole proteomes were sliced and treated with iodoacetamide and then trypsin as previously recommended by (Hartmann et al., 2014). Briefly, each band was destained with ultra-pure water, reduced with dithiothreitol, treated with iodoacetamide, and then proteolyzed with Trypsin Gold Mass Spectrometry Grade (Promega) in the presence of 0.01% ProteaseMAX surfactant (Promega). Peptides were immediately subjected to tandem mass spectrometry as previously recommended to avoid methionine oxidation (Madeira et al., 2017). The resulting peptide mixtures were analyzed in a data-dependent mode with a Q-Exactive HF tandem mass spectrometer (Thermo) coupled on line to an Ultimate 3000 chromatography system chromatography (Thermo) essentially as previously described (Klein et al., 2016). A volume of 10 μL of each peptide sample was injected, first desalted with a reverse-phase Acclaim PepMap 100 C18 (5 μm, 100 Å, 5 mm x 300 μm i.d., Thermo) precolumn and then separated at a flow rate of 0.2 μL per min with a nanoscale Acclaim PepMap 100 C18 (3 μm, 100 Å, 500 mm x 300 μm i.d., Thermo) column using a 150 min gradient from 2.5 *%* to 25 *%* of CH3CN, 0.1% formic acid, followed by a 30 min gradient from 25% to 40% of CH3CN, 0.1% formic acid. Mass determination of peptides was done at a resolution of 60,000. Peptides were then selected for fragmentation according to a Top20 method with a dynamic exclusion of 10 sec. MS/MS mass spectra were acquired with an AGC target set at 1.7 10^5^ on peptides with 2 or 3 positive charges, an isolation window set at 1.6 *m/z,* and a resolution of 15,000.

### MS/MS spectrum assignment, peptide validation and protein identification

Peak lists were automatically generated from raw datasets with Proteome Discoverer 1.4.1 (Thermo) and an in-house script with the following options: minimum mass (400), maximum mass (5,000), grouping tolerance (0), intermediate scans (0) and threshold (1,000). The resulting .mgf files were queried with the Mascot software version 2.5.1 (Matrix Science) against the *R. sphaeroides* 241 annotated genome database with the following parameters: full-trypsin specificity, up to 2 missed cleavages allowed, static modification of carbamidomethylated cysteine, variable oxidation of methionine, variable deamidation of asparagine and glutamine, mass tolerance of 5 ppm on parent ions and mass tolerance on MS/MS of 0.02 Da. The decoy search option of Mascot was activated for estimating the false discovery rate (FDR) that was below 1%. Peptide matches with a MASCOT peptide score below a p value of 0.05 were considered. Proteins were validated when at least two different peptides were detected. The FDR for proteins was below 1% as estimated with the MASCOT reverse database decoy search option.

### Ice logo analysis

Ice logoanalysis were performed using the IceLogo server (http://iomics.ugent.be/icelogoserver/index.html) (Colaert et al., 2009).

